# Population responses to environmental stochasticity are primarily driven by survival-reproduction trade-offs and mediated by aridity

**DOI:** 10.1101/2024.07.24.604949

**Authors:** Gabriel Silva Santos, Xianyu Yang, Samuel J L Gascoigne, Aldo Compagnoni, André T.C. Dias, Shripad Tuljapurkar, Maja Kajin, Roberto Salguero-Gómez

## Abstract

Forecasting responses of natural populations to increasingly stochastic environments is a major challenge in Ecology and Conservation Biology. We now know that populations can modulate how their vital rates (*e*.*g*., survival, reproduction) change through time to minimise the negative impacts of environmental stochasticity. However, despite the important analytical and theoretical advances that have led to this knowledge, we still do not know (1) how much this ability of natural populations to buffer against environmental stochasticity can vary in nature, nor (2) the drivers of these strategies, with likely candidates including the environmental regimes themselves, as well as the life history traits and phylogenetic ancestry of the species of interest. To address these questions, we parameterised a Bayesian generalised linear mixed model with high-resolution vital rate data from 134 natural populations across 89 species of plants and animals. We show that population responses to environmental stochasticity vary three orders of magnitude along a ‘demographic buffering continuum’. Furthermore, the position of a given population along said continuum is predicted by a survival*-*reproduction trade-off and by the degree of aridity the population experiences. Our findings open a promising avenue of research to improve ecological forecasts and management of natural populations in the Anthropocene.

## 1. Introduction

The long-term viability of natural populations is largely determined by environmental stochasticity^1,2^. Climate change projections anticipate a global increase in not only the mean of key abiotic drivers (*e*.*g*., temperature, precipitation), but also their temporal variance^3^. Crucially, increased environmental stochasticity has already brought about important eco-evolutionary challenges for the performance and viability of natural populations^2,4^, and has been identified as a major driver of biodiversity loss worldwide^5^. As such, understanding whether and how populations minimise the expected negative effects of environmental stochasticity has become a primary mission of Ecology, Evolution, and Conservation Biology^6^.

Half a century of research examining how species’ life history strategies are shaped by natural selection to cope with environmental stochasticity have led to the establishment of two major axes of life history variation: the fast-slow continuum^7,8^ and the reproductive strategies continuum^9–11^. The fast-slow continuum ranks organisms according to a development-survival trade-off from fast growing, short-lived organisms to slow growing, long-lived ones^7^. In contrast, the reproductive strategy continuum, which is orthogonal to the fast-slow continuum^12–14^, categorises organisms based on the length of their reproductive window, from single (*i*.*e*., semelparous) to multiple reproductive bouts (*i*.*e*., iteroparous)^9,15^. Recent advances in biodemography have linked species at the slow-end of the fast-slow continuum to a greater capacity to buffer the negative impacts of environmental variation mediated by natural selection^16– 18^. This so-called demographic buffering^17,19^ is accomplished by reducing the temporal variance of those vital rates (*e*.*g*., survival, growth, reproduction) that contribute most to stochastic population growth rate (*λ*_*s*_). However, the demographic buffering hypothesis^17,19^ does not account for the various shapes of environment-vital rate reaction norms^20–22^. For instance, following Jensen’s inequality^23^, convex (∪-shaped) environment-vital rate reaction norms can result in a positive effect of vital rate variance on *λ*_*s*_, whereas linear or concave (∩-shaped) reaction norms lead to a negative effect^21,24^. Importantly, there are limits to the amount of variance that a vital rate can exhibit without driving a population to local extinction^25^. Altogether, these complementary parts to the demographic buffering hypothesis remain a major challenge to link life history continua to populations response to environmental stochasticity in the context of climatic change.

Here, we focus on a continuum of vital rate temporal variance to examine how environmental regimes, key life history traits, and evolutionary history predict the extent to which natural populations of multicellular organisms can buffer against environmental stochasticity^26^. Specifically, we parameterise a phylogenetically-corrected Bayesian Generalised Linear Mixed Model (GLMM) with high-resolution vital rate data from 134 natural populations across 11 animal and 78 plant species to test the following hypothesis: (*H*_*1a*_) Plant and animal populations’ responses to stochastic environments regarding vital rate variance vary widely, but largely overlap along a continuum of demographic buffering where, on one extreme, temporal variance of governing vital rates is constrained (*i*.*e*. more buffered, Fig. 1), while on the other extreme, not at all (less buffered). Additionally, (*H*_*1b*_) the position of species’ populations along the demographic buffering continuum is not predicted by the fast-slow and reproductive strategies continua. Although a study using 36 populations of different species found that longevity (a life history trait aligned with the fast-slow continuum^10–12,14^) is a good proxy of demographic buffering^18^, another study with a greater number of populations and that explicitly accounted for their evolutionary history provided opposing evidence^27^. However, none of these studies included environmental metrics to explicitly account for the actual environmental stochasticity faced by the studied population. This key omission overlooks the potential roles of reproductive strategies, phylogenetic relationships, and environmental regimes on population responses to environmental stochasticity; (*H*_*2*_) phylogenetic ancestry will play a stronger role in determining demographic buffering capacities in animals than in plants because morphology and behaviour, strong predictors of a species’ demography^28,29^, are much more conserved across animals than in plants^30,31^; and (*H*_*3*_) the extent to which a natural population displays demographic buffering will depend on the environmental regime that it has been exposed to, with higher stochasticity in precipitation and temperature contributing to greater variances in vital rates, which will place populations toward a less buffered end of said continuum.

**Figure 1.**
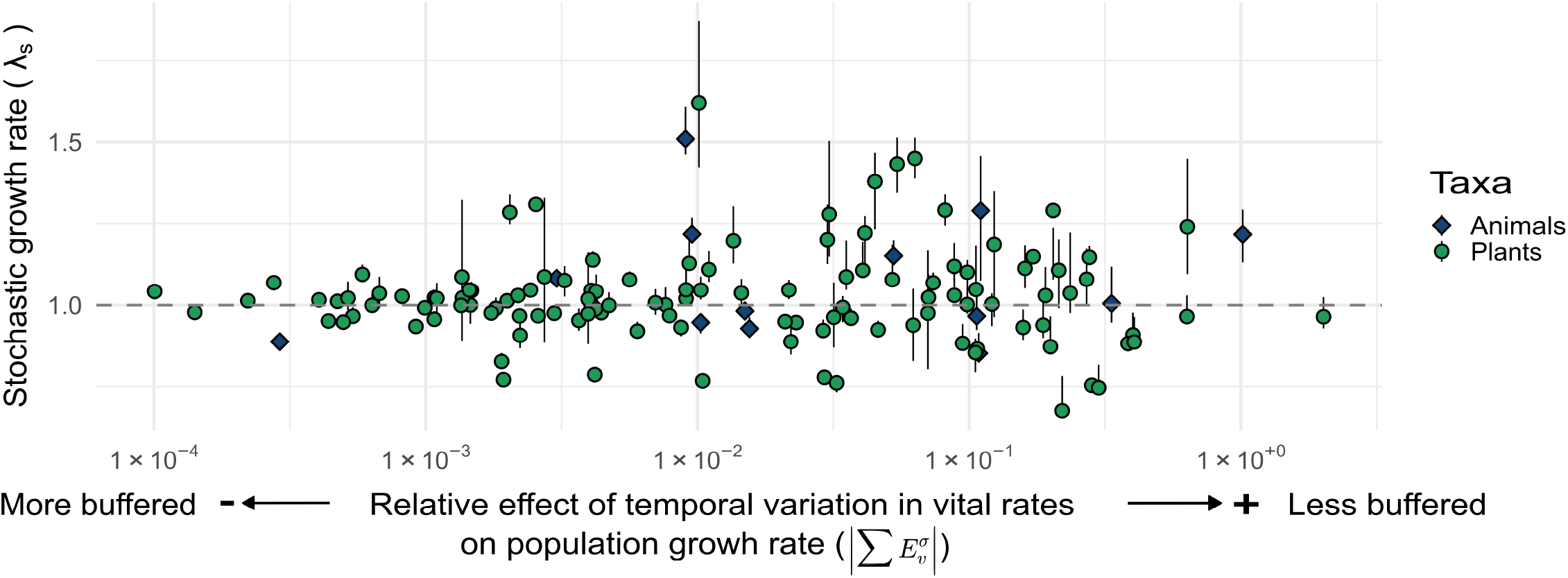
Plant (green) and animal (blue) range along a continuum of demographic buffering against effect of temporal variation in vital rates. Said continuum is quantified by the sum of stochastic elasticities within respect to the variance 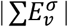 in the vital rates (*i*.*e*., survival, individual-level growth, individual-level shrinkage, reproduction, and clonality). Values of 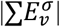 represent the extent to which changes in the temporal variation of a given vital rate affect the stochastic population growth rate (*λ*_*s*_) in 134 natural populations (13 animals and 121 plant species) retrieved from the COMADRE (version 4.23.3.1) and COMPADRE (v. 6.23.5.0) databases. At the buffering-end of the continuum (right; 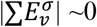), vital rates either vary less or the observed variation has little to negligible effect on *λ*_*s*_. At the not buffered end (left; 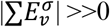), the observed variation on vital rates strongly shapes *λ*_*s*_ values.

To test our hypotheses, we first established the existence of a continuum of vital rate variance from high to low demographic buffering. This continuum emerges when examining how the observed temporal variance of vital rates has shaped the long-term population performance of each natural population. To do so, we use the sum of stochastic elasticity of the stochastic population growth rate *λ*_*s*_ with respect to vital rate temporal variance, 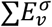. This variable was calculated from a subset of matrix population models from COMPADRE^32^ and COMADRE^33^ databases that depict the demography of populations under natural environmental conditions (see Materials and Methods for further details on the data selection criteria). Absolute values of 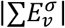 close to 0 correspond with small effects of the temporal variation of vital rates on population growth rate *λ*_*s*_(*i*.*e*., high demographic buffering), while 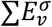 away from 0 represents a more tangible effect on *λ*_*s*_(low demographic buffering). Next, we also quantified the responses of our 134 natural populations to environmental stochasticity in the context of their positions along multiple axes of variation: the fast-slow and reproductive continua, and environmental stochasticity, using Principal Component Analyses (PCAs).

## 2. Results

Our 134 examined plant and animal populations unveil a continuum of variance in vital rates that is highly overlapping between the Kingdoms, as quantified by 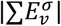. In fact, we find no evident distinction in the range of demographic buffering responses to environmental stochasticity between plants and animals (*t*-test = 0.701, *p* = 0.495; Figure 1). These findings provide support for our hypothesis *H*_*1a*_, that the responses of natural populations to environmental stochasticity might be described along a continuum from strongly buffering to not at all, independent of kingdom. For animals, the variation in demographic buffering ranges from rather low values, such as 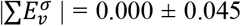 (SD) in the Red gorgonian (*Paramuricea clavata*) to 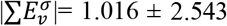 in the Woolly sculpin (*Clinocottus analis*). For plants, the demographic buffering ranges again from low values, like 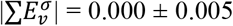 for the Pyrenean violet (*Ramonda myconi*) to an even higher maximum value than for animals: 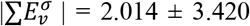 in the Mexican hat (*Ratibida columnifera*). For both kingdoms, the position of populations along the demographic buffering continuum is mainly determined by the temporal variance of survival (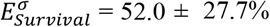; Figure 2), followed by individual-level growth 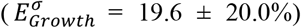 and reproduction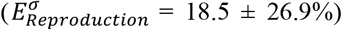, with minor contributions from individual-level shrinkage 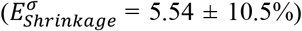 and clonality (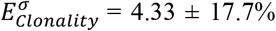; Figure 2 and Table S1).

**Figure 2.**
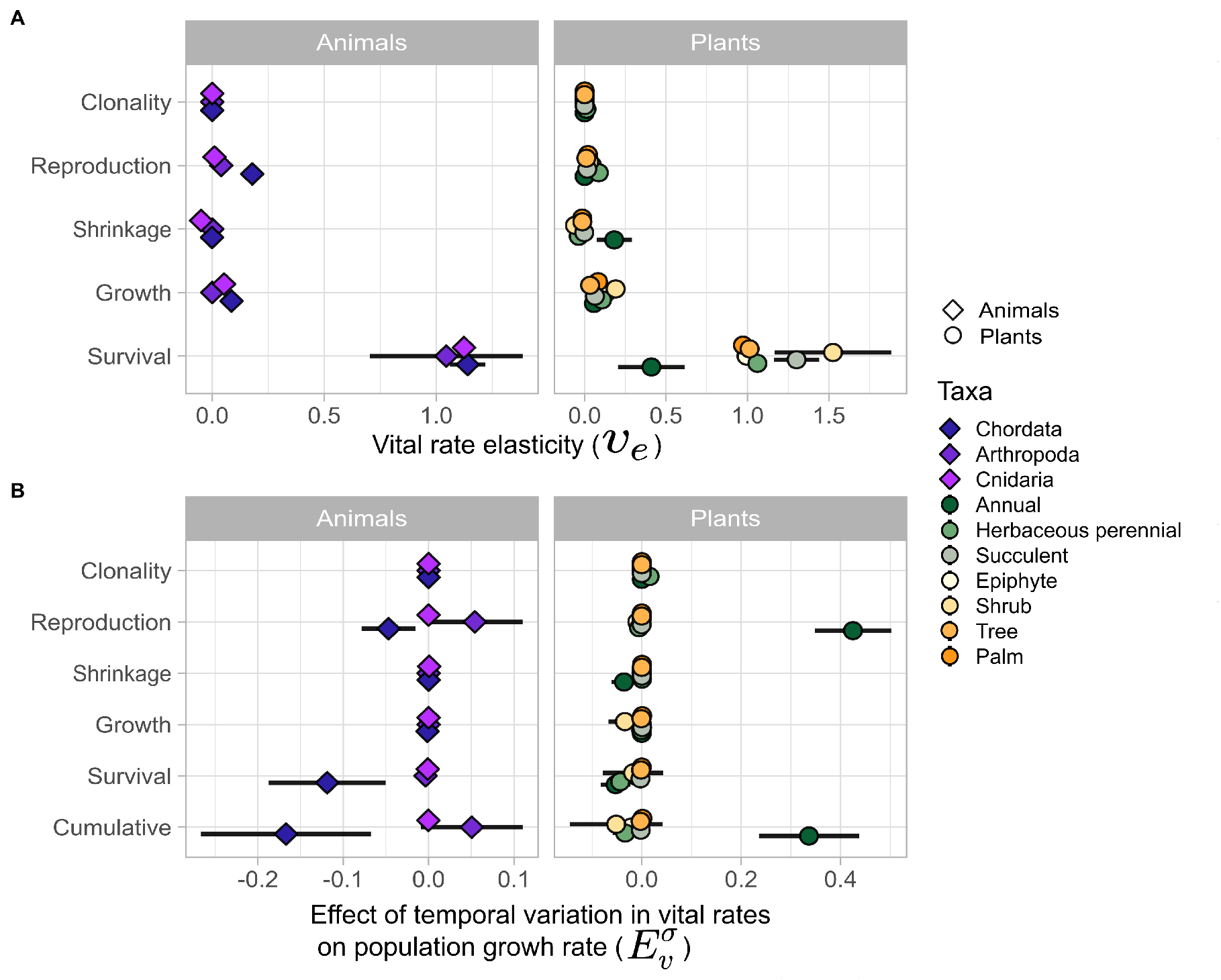
For most plants and animals, the temporal variance in survival is the main driver of the responses of natural populations to environmental stochasticity, followed by individual-level growth and reproduction. **A**. The relative importance of each vital rate (elasticities, *E*_*v*_ ) to population growth rate, *λ*_*s*_ (Mean: circle; S.E.: bars). **B**. Stochastic elasticity of *λ*_*s*_ with respect to the vital rate temporal variance 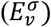 of the vital rates: survival, growth, shrinkage, reproduction, and clonality. “Cumulative” refers the cumulative variance effect of all vital rates on *λ*_*s*_, 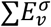. Plants and animals are represented by different colours and shapes, with colours being subset based on taxonomic Order for animals and life-forms for plants. Overall, 134 populations of plants (n=121) and animals (n=13) were included.

Our analyses provide support for *H*_*1b*_ in plants, but not in animals. Indeed, the position of plant populations along the demographic buffering continuum, as quantified by 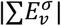, is predicted by the fast-slow and reproductive strategies continua (Figure 3). However, in plants, the reproductive strategies continuum is only informative when phylogenetic ancestry is not considered (Figure 4). Because of the lack of support for *H*_*1b*_ in animals, we focus our results for plants populations hereafter, but both results are presented (see Fig. 3 and 4). Thus, considering plants populations, our model without phylogenetic corrections, the interaction between the fast-slow and reproductive strategy continua (the first two principal component axes of life histories, *PC*1_*LH*_ : *PC*2_*LH*_, respectively; Fig. S1) determines the position of populations along the demographic buffering continuum (non-phylogenetic corrected MCMCglmm 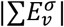, 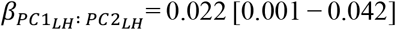; posterior mean and 95% credible interval). However, neither main effects of *PC*1_*LH*_ (fast-slow continuum) nor *PC*2_*LH*_ (reproductive continuum) are significant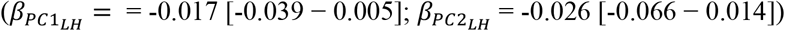. More importantly, after controlling for phylogenetic relationships in our data, only the fast-slow continuum remains an important predictor of 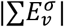 in plants (phylogenetically corrected MCMCglmm 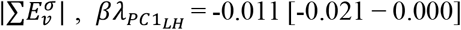; Figure 3), with no significance for animals. Thus, in agreement with our hypothesis *H*_*1b*_, there is no link between the reproductive strategy continuum and the contributions of reproduction to the demographic buffering capacity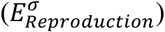, independent of phylogenetic corrections 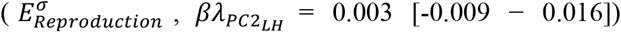 or not (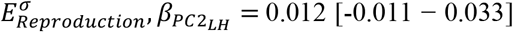Figure 4). The lack of relationship between the reproductive strategy continuum and demographic buffering continuum may reflect: (i) the reduced impact temporal variation in reproduction has on *λ*_*s*_(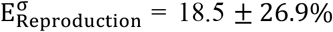; Figure 2 and Table S1), and/or (ii) the strong phylogenetic signal in the contribution of reproduction to 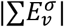, at least for plants (Pagel’s *λ*_*reproduction*_ = 0.971± 0.011; Figure 4). This result suggests that the contribution of reproduction to the populations’ responses to environmental stochasticity is better predicted by phylogenetic ancestry than by life history traits or environmental conditions.

**Figure 3.**
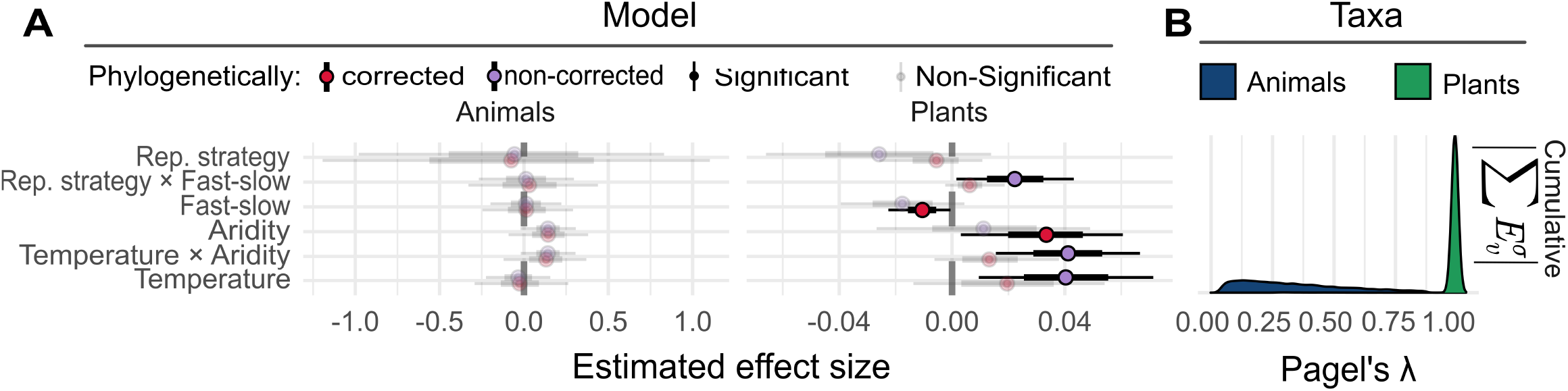
The importance of life history traits and environmental stochasticity on the demographic buffering capacity of natural populations depends strongly on phylogenetic relationships. **A**. Predictive power of the life history principal component (PC) axes and environmental stochasticity PC on the positioning of species’ populations along the buffering continuum, measured as the 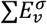. The shown posteriors were obtained via MCMCglmm models with (red) and without (purple) phylogenetic corrections. The fast-slow continuum and reproductive strategies continuum is represented by *PC*1_*LH*_ and *PC*2_*LH*_ (Figure S1), respectively; whereas environmental stochasticity in temperature and aridity represents the variation in the environment experienced by the examined populations as described by *PC*1_*Env*_ and *PC*2_*Env*_ (Figure S2). Significant relationships are shown in black, non-significant are blurred. **B**. Distribution of our estimates of phylogenetic signals (Pagel’s *λ*) for animals (blue) and plants (green) from our phylogenetic MCMCglmm model, evidencing the importance of phylogenetic inertia in determining how life history strategies and environmental regimes shape population responses to environmental stochasticity, 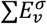.

**Figure 4.**
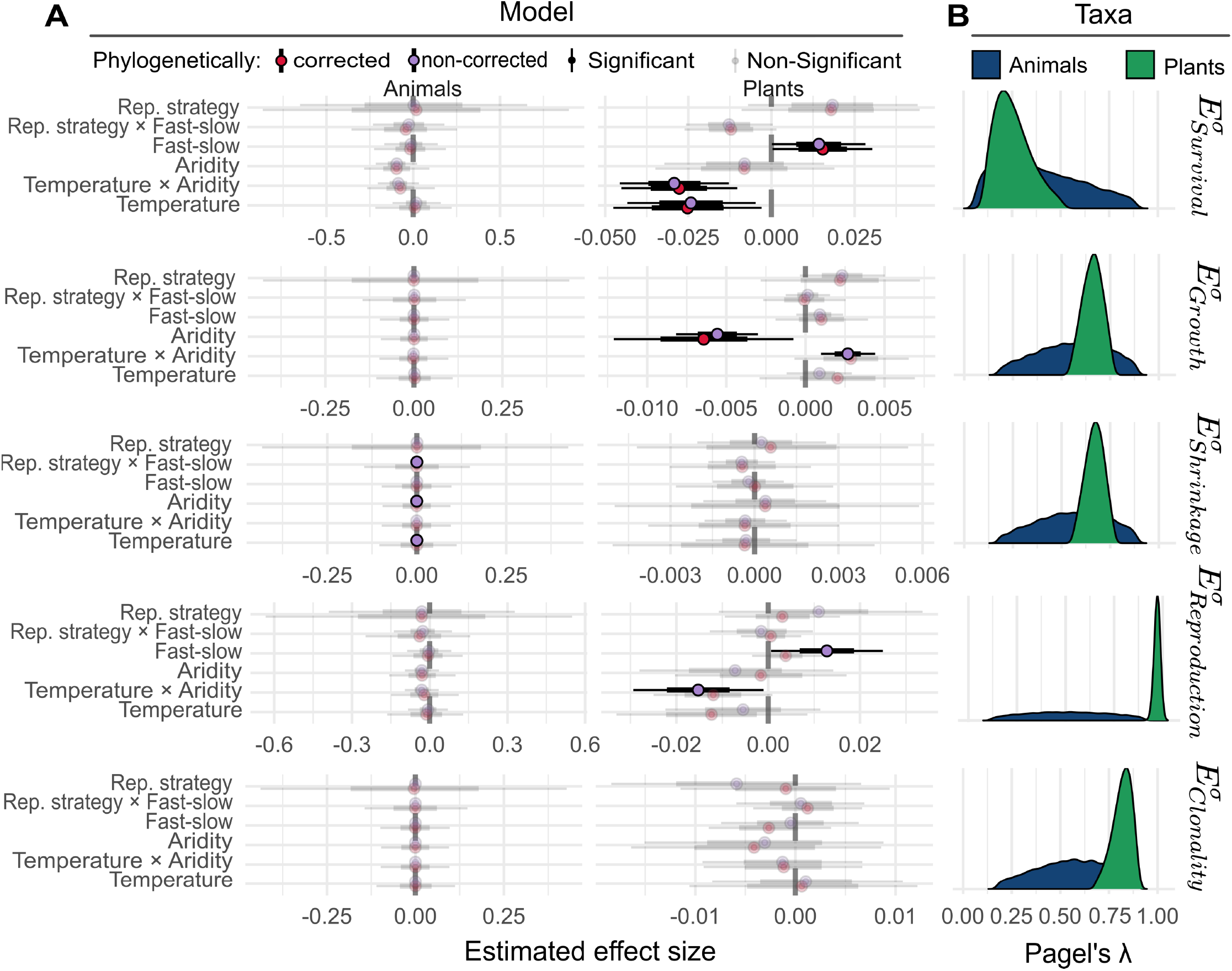
Life history traits and environmental stochasticity have variable contributions on the effects of the temporal variation in vital rates, *v*, on the stochastic population growth rate, *λ*_*s*_. **A**. Predictive power of the life history principal component (PC) axes and environmental stochasticity PC on the positioning of species’ populations along the demographic buffering continuum, measured as 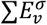. The fast-slow continuum and reproductive strategies continuum is represented by *PC*1_*LH*_ and *PC*2_*LH*_ (Figure S1), respectively; whereas environmental stochasticity in temperature and aridity represents the variation in the environment experienced by the examined populations as described by *PC*1_*Env*_ and *PC*2_*Env*_ (Figure S2). **B**. Distribution of our estimates of phylogenetic signals (Pagel’s *λ*) for animals (blue) and plants (green) from our phylogenetic MCMCglmm model, evidencing a variable importance of phylogenetic inertia in determining how life history strategies and environmental regimes shape population responses to stochasticity,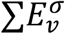.

The aforementioned discrepancies between phylogenetically and non-phylogenetically corrected models support the idea that evolutionary history plays an important role in the placement of populations along the demographic buffering continuum. However, our results do not support *H*_*2*_, that phylogenetic ancestry would play a stronger role in animals than in plants. Instead, the values of 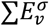 along the demographic buffering continuum for plants raise a very strong phylogenetic signal (Pagel’s *λ* = 0.967 ± 0.138), while animals have a relatively weak one (Pagel’s *λ* = 0.345 ± 0.230). However, the phylogenetic signal of the contributions of the underlying vital rates to 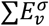 is variable. For instance, survival has a rather weak phylogenetic signal for both plants (Pagel’s *λ*_*survival*_ = 0.259 ± 0.106; mean ± SD) and animals (Pagel’s *λ*_*survival*_ = 0.399 ± 0.229; Figure 4). A potential reason to explain why the plant demographic buffering continuum has a higher phylogenetic signal than animals, even if it does not hold for survival, may be the greater sampled size in plants, which includes some species represented by multiple populations in our study (Table S2). Consequently, more conservative values of 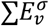 between populations of the same species compared to other species might be promoting a higher phylogenetic signal in plants. The same relationship might in reality apply to natural populations of animals, an aspect that we might be able to unveil as the discipline continues to accrue longer, high-quality vital rate data^35^.

Our phylogenetically and non-phylogenetically corrected models both support *H*_*3*_, that populations located in more stochastic environments are positioned towards the low-buffering end 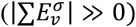 of the demographic buffering continuum (Fig. 1). In our 134 natural populations, temporal variation in aridity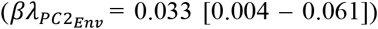, more so than in temperature (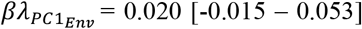; Figure 3), act as the main determinant of the placement of populations along the demographic buffering continuum. Moreover, the temporal variation in aridity 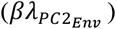 outweighs over three-fold the predictive power of the fast-slow continuum 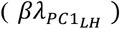 (posteriors 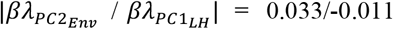). However, the influence of environmental stochasticity on population responses depends on the vital rate under consideration. For example, aridity is the main determinant of the contribution of individual growth to population responses to environmental stochasticity 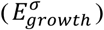, with more stochasticity in aridity being associated with lower demographic buffering, at least for plants (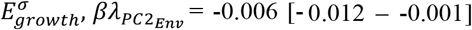; Figure 4). In contrast, temperature and aridity act together to determine the importance of survival 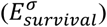 on 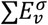 values of plants (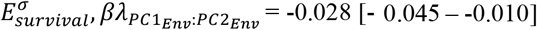; Figure 4). These findings likely reflect the greater dependence of plants, as sessile organisms, on the capacity to survive and grow under heatwave-induced drought events^36^ compared to the – mostly mobile – animal species examined here.

## 3. Discussion

The fast-slow and reproductive strategies continua of life history traits have long dominated predictions of species adaptation to environmental conditions^37,38^. However, these axes have proven limited in predicting population responses to environmental stochasticity^39–41^. Over the past two decades, a novel framework to examine population responses to environmental stochasticity has emerged based on the idea that natural selection must canalise the temporal variation in vital rates to buffer against the expected negative impacts of environmental stochasticity^19,42^, a prediction referred to as the ‘demographic buffering hypothesis’^19,42^. Despite significant progress in the pertinent theory^17^ and praxis^43^, we still lack a solid understanding of the drivers of population responses to environmental stochasticity, their relationship to the well-known fast-slow and reproductive strategies continua, and the role of evolutionary history and environmental regimes^17,27^. Here, we provide a critical assessment of this gap in knowledge and report several key findings: (i) Plant and animal populations fall along a single axis, with significant overlap, that quantifies their responses to environmental stochasticity, from more to less demographically buffered; (ii) This response can be defined by species’ life history (*i*.*e*., fast-slow and reproductive strategies), phylogenetic relationships, and environmental conditions; however, these relationships are complex; (iii) When phylogenetic relationships are taken into account, the fast-slow continuum – but not the reproductive strategies continuum – remains as an important predictor of species response to environmental stochasticity; (iv) Yet, the predictive power of the fast-slow continuum on the ability of a population to buffer against the environment is outweighed by a key aspect of environmental stochasticity: variation in aridity. Finally, (v) species’ responses to environmental stochasticity are strongly cemented on evolutionary history, but, contrary predictions^31^, the phylogenetic signal is stronger in plants than in animals.

In our study, environmental stochasticity outweighs the role of life history traits in predicting the buffering capacities of natural populations. This effect is explained by the larger influence of survival on species’ buffering capacity, which we also characterise as the vital rate with lowest phylogenetic conservatism in our study, well above growth, shrinkage, and reproduction. As consequence, the position of a given population along the demographic buffering continuum reflects its capacity to buffer variation in survival, a result that supports the well-known challenge of predicting survival of animals and plants in a stochastic environment^36,41^. Here, we highlight three major challenges to use evolutionary history and life history traits to predict plant and animal survival in a stochastic environment. First, organisms may possess life history traits that were once advantageous in the ancestral environments but that, given the fast changing climate^3^, are no longer beneficial. For instance, some species have evolved specific windows for recruitment or migration that were adaptive in environments with predictable fluctuations, but are maladaptive in current/future unpredictable environments^44^. Second, individual responses to environmental stochasticity are highly heterogeneous and tend to be determined by their capacities to obtain and conserve energy rather than their evolutionary life history^41^. Third, covariation between physiological and morphological traits can amplify or dampen the impact of environmental stochasticity on individual performance^36,45,46^. Consequently, environmental stochasticity may result in nonlinear responses that pose important challenges for ecological forecast^21,47^. Together, these challenges likely weaken the predictive power of the fast-slow and reproductive strategies axes, which do not explicitly consider demographic variation. As such, we argue that these life history axes reflect how species have evolved to cope with past environmental stochasticity rather than current or future pressures populations – are likely to – face. On the other hand, these complex responses exemplify the numerous evolutionary processes that can influence temporal variation in vital rates, upon which the demographic buffering hypothesis was developed^22,48^. The existence of a demographic buffering continuum that quantifies the temporal variance in vital rates, as we report here, is a promising tool to generate new insights regarding which and how populations respond to environmental stochasticity. It is worth noting that, for plants and animals, populations with higher long-term performance (*λ*_*s*_) are located at intermediary levels of temporal variance in their vital rates, suggesting that some variance is potentially beneficial, but too much may not be.

Despite being subject to stronger phylogenetic conservatism, the contribution of reproduction to population response to environmental stochasticity is relatively low compared to other vital rates. This finding contrasts with the high influence but low phylogenetic conservatism of survival. Together, these findings call for more research exploring how survival-reproduction trade-offs will shape population performance in the Anthropocene. For instance, recent studies have shown how reductions in survival might be compensated by increases in reproduction, a phenomenon known as demographic compensation^49,50^. Although a growing body of literature has supported the widespread existence of demographic compensation among natural populations^50,51^, their capacity to compensate against the negative impact of environmental stochasticity remains poorly understood^49^. Given our finding of low contribution but high predictable influence of reproduction on population responses to environmental stochasticity, one could expect limited contributions of reproduction in this compensation process as well as great predictability of their compensation capacity. Investigating such a questions is particularly key to predict population responses to climate change^52^, particularly under uncertain increasing in environmental stochasticity^3^. Additionally, we suggest that, because the effects of reproduction on population responses to environmental stochasticity are highly predictable by evolutionary history, there is an opportunity to improve the parameterisation and accuracy of demographic models in population viability analyses^53^. This opportunity includes models examining early impacts of exploitation, such seed harvesting^54^. When survival (most important and less predictable vital rate) cannot be fitted in the model parameterization, reducing uncertainty in other vital rates such reproduction is imperative^53^. Our findings show promising avenue of research in this direction.

It is critical to recognise that our framework, using the sum of stochastic elasticities of the stochastic population growth rate *λ*_*s*_with respect to vital rate temporal variance, 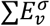, has two areas for improvement. First, the theoretical aspect related to the buffering hypothesis and its alternative hypothesis – the demographic lability hypothesis^19,21,48^ – remains poorly understood. This gap of knowledge hampers a direct link between to the demographic buffering continuum of variance explored in this study. The demographic buffering and lability hypotheses have been understood in terms of underlying evolutionary mechanisms such as canalising by stabilising selection resulting in buffering, or boosting the temporal variance by disruptive selection in lability^19,21,48^. Ultimately, these evolutionary processes might result in the observed patterns of vital rate variance, from constraints (if buffered) to boosting (if labile), which have been the analytical focus of the attempts to test these hypotheses in recent years^19,27,43^. However, such underlying evolutionary mechanisms of buffering and lability have only been inferred, instead of being directly assessed^48^. So far, the only two attempts to explicitly identify and quantify the underlying evolutionary mechanisms behind demographic buffering and lability (nonlinear indexes^22^ and second-order derivatives^48^) have not considered a critical component: the observed temporal variation and its impact on population growth rate. Our approach captures only the temporal variation and its impact on population performance, which represents their overall demographic buffering capacity, but not the underlying evolutionary mechanisms (*i*.*e*., stabilizing *vs*. disruptive selection) driving this temporal variation^48^. Thus, while our framework is not the final stop of this important line of enquiry, our framework provides a unique set of opportunities to complement recent evolutionary metrics such as nonlinearity indexes^22^ and second-order derivatives^48^. Further studies may integrate such metrics into the present framework to link the forces of natural selection and their effects on the current and future performance of natural populations.

A second area for improvement is the suboptimal representation of long-term demographic studies across taxonomies and geographies required for comparative demographic studies. The existing databases we use here, though state-of-the-art in regards to number of demographic studies, still remains biased toward perennial species in the Plant Kingdom, and mammals and birds in the Animal Kingdom, as well as towards temperate habitats across both kingdoms^55,56^. Moreover, only a rather small number of animal species currently have a sufficiently long time-series of demographic models to be included in our analytical pipeline, given our strict selection criteria (see Methods). Filling the demographic gap across continental scales, particularly South America and Africa, remains a pressing issue to gain a holistic understand of how natural populations respond to environmental stochasticity^55,56^. High-resolution demographic data, however, take by definition a long time to be produced, which makes filling this gap an enduring commitment by population ecologists in the coming decades.

Ecology and conservation biology will improve the ecological forecasting of population responses to environmental conditions by considering a more flexible life history axis that explicitly accounts for the capacity of natural population to respond to environmental stochasticity: the *demographic buffering continuum*. Indeed, this continuum explicitly quantifies how populations respond to environmental stochasticity by evaluating their effects on long-term population viability in real environments. By developing and applying the demographic buffering continuum here, we succeeded in quantifying the effect of environmental stochasticity on population responses and linking these responses to life history traits and their evolutionary history. Mostly important, our approach allows us to show that plants and animals share commonalities in how they respond to environmental stochasticity, suggesting that distinguishing multicellular organisms by taxonomic boundaries may hinder our understanding of the responses of natural populations to environmental stochasticity^57^. Together, our approach and findings highlight a promising way to better parameterise population models and improve population forecasts in the Anthropocene.

## 4. Materials and Methods

To test our hypotheses, we developed a Bayesian Generalised Linear Mixed Model (GLMM) with and without phylogenetic corrections (see Eq. 1). Doing so allows us to test the robustness of our results to the role of shared ancestry in our comparative analyses. Having established a demographic buffering continuum of variance (Fig. 1), we then performed separate models to better understand the importance of five vital rates (survival, growth, shrinkage, reproduction, and clonality) in determining the position of species’ populations along the said continuum. Briefly, to position those populations on a multivariate space and test for the existence of a demographic buffering continuum, we derived life history traits from Matrix Population Models (MPMs;^58,59^). To examine the role of environmental regimes on demographic buffering, we extracted environmental data from CHELSAcruts^60^ during the period when each population was studied. Data collection, model parametrisation, and the general analytical approach are detailed below.

### Demographic data and species responses to environmental stochasticity

To assess the roles of vital rates in shaping stochastic population growth rates (*λ*_*s*_), we used time series of MPMs from natural populations. An MPM is a discrete-state mathematical representation of the life cycle of a species, typically represented by a matrix ***A***. Each matrix element *a*_*ij*_ in a ***A*** represents the contribution of a current (st)age to the next (st)age via survival, growth (development/progression), shrinkage/retrogression, and reproduction of individuals^58,61^. These MPMs can be decomposed into submatrices representing the different processes of the life cycle: the ***U*** submatrix describes survival-dependent transitions (*e*.*g*., progression, retrogression), while the ***F*** and ***C*** submatrices describe sexual and clonal reproductions, respectively^59^. The MPMs and their decomposed submatrices used in this study were selected from the COMPADRE Plant Matrix Database v6.23.5.0^32^, and the COMADRE Animal Database v4.23.3.1^33^, which contain 792 plant species with 8,994 MPM and 429 animal species with 3,488 MPMs, respectively.

Population responses to environmental stochasticity were analysed by estimating the overall effect of temporal variation in vital rates on the stochastic population growth rate *λ*_*s*_. Population responses were assessed via the well-established method of the sum of stochastic elasticities within respect to variance, 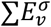^48,61,62^. Briefly, this approach estimates the extent to which small changes in the mean and variance of a given vital rate *v* (or matrix element, *a*_*ij*_) affects *λ*_*s*_. In mathematical terms, stochastic elasticities are the partial derivative of the vital rates *v* (or matrix elements) 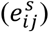 over the MPM time series weighted by the relative contribution to *λ*_*s*_, and are typically expressed as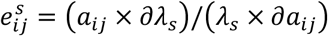. The overall contribution of each vital rate 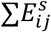 to *λ*_*s*_ is always 1, but said overall contribution can be partitioned into the effect of perturbing the mean (*μ*) values of vital rate 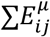 or its variance (*σ*), 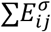, such that 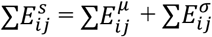. The variance component 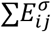 ranges between 0 to -∞, where values close to zero means that vital rate temporal variation has a nearly negligible impact on *λ*_*s*_^48^.

For illustration, consider a vital rate *v* tracked over time in a population. Because environmental conditions are not static, some degree of change in these vital rates may be expected between the intervals *t* →*t*+1, *t*+1 → *t*+2, *etc*. The differences in the values *v* and their effect on *λ*_*s*_each time step represents the population’s response to the environment during the said time interval (or due to lag effects; see^63^). The relative impact of each vital rate *v* on *λ*_*s*_can be readily assessed by its associate deterministic elasticity value, and the cumulative sum of these stochastic elasticities with respect to the temporal variance in *v* can thus be interpreted as the overall demographic impact on *λ*_*s*_^61,62^.

Importantly, while previous works have developed this approach for matrix elements^34,48,62,64^, here we extend the method to evaluate the effects of the underlying components of matrix elements – vital rates, *v* – thus offering higher resolution and allowing evaluating the impact of survival independent of other survival-dependent processes such as growth and shrinkage^65^. Unlike matrix elements, elasticities in the underlying vital rates might be negative^65^, thus the sum of stochastic elasticity of *λ*_*s*_with respect to vital rate temporal variance for the underlying vital rates ranges from -∞ to +∞. However, the main pattern persists, with 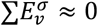 representing more demographically buffered populations, and 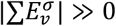 representing populations where temporal variation of vital rates strongly affects *λ*_*s*_, either negatively or positively. As such, our approach explicitly accounts for the way that a given environmental regime may impact vital rates and how these vital rates, in turn, shape the overall performance of the population^61,62^. In our study, vital rates were retrieved from MPMs organised into three submatrices. ***U, F*** and ***C***^59,65^. Specifically, survival, growth, and shrinkage were obtained from submatrix ***U***, whereas sexual reproduction and clonal reproduction were estimated from the submatrices ***F*** and ***C***, respectively. To do so, we used the family function *vr_* (*e*.*g*., *vr_growth, vr_shrinkage)* in the *Rage* R package^59^. Having access to already decomposed submatrices in COMPADRE and COMADRE, we explored the proportional contribution of each vital rate (*e*.*g*.,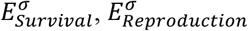) to 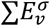, represented by 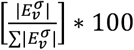.

The variable 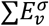 estimates population responses to environmental stochasticity by recognising multiple evolutionary and physiological processes. For instance, physiological constraints might limit variation in reproduction^66^ as well as canalising selection^67^, both evolutionary processes pushing populations to the buffering end^48^. Meanwhile, disruptive selection and high phenotypic plasticity might support adaptative variation, thus pushing populations to less-buffered end^21,47^. Furthermore, high values of 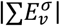 indicate heightened responsiveness to environmental stochasticity, regardless of whether this responsiveness is favoured by natural selection^62^. Despite limited inference, this approach overcomes existing approaches by: (1) considering the environmental stochasticity experienced by populations not explicitly accounted in the nonlinearity index, which is calculated from their mean MPMs^22^; and (2) not being affected by constrained variation existing near the extreme values of vital rates that have puzzled the correlative approaches between vital rate variance and sensitivities^17,19,27^.

### Demographic data selection and phylogenetic relationships

To ensure comparability across and within species, we selected MPMs from COMPADRE and COMADRE that fulfilled a series of carefully planned selection criteria to ensure comparability and robust results: (1) only natural populations (*e*.*g*., no zoo data) to explicitly link vital rates to their natural environments; (2) studies containing MPMs with at least three annual contiguous censuses to capture environmental stochasticity in the vital rates; (3) irreducible, primitive, and ergodic MPMs to ensure that each MPM represents a complete life cycle^68^; (4) MPMs with separated ***U, F***, and ***C*** submatrices that allow us to disentangle the effects of vital rates on the overall population growth rate obtained from the matrix ***A***; (5) populations with known GPS coordinates, so we could match the environmental data during the study period (see below) to vital rate variation; and (6) MPMs whose species are present in a well-resolved phylogeny to test the role of phylogenetic inertia on demographic buffering continuum (*H*_*3*_). Finally, we removed five populations identified as outliers (see *Life history traits, life history axes, and the life history PCA*). These selection criteria yielded a total of 889 MPMs across 134 populations of 89 species, of which 121 populations belonged to 78 plant species, and 13 populations to 11 animal species.

To account for the evolutionary history in our models, we used the phylogenetic trees available in the MOSAIC database^69^. The trees are continuously updated from the EOL project^70^ and comprise most of the species available in COMPADRE and COMADRE. In them, polytomies were resolved using the *multi2di* function from *ape* R package^71^. Further details regarding the construction of the phylogenetic trees are found in Salguero-Gómez et al. ^12^ for plants and Healy et al.^14^ for animals.

### Life history traits and axes of variation

The position of each species’ populations along the fast-slow and reproductive strategy continua was used as covariates to test *H*_*1b*_ (below). To assess the position of each of the 134 populations along these two axes, we performed principal component analyses of the six examined life history traits. Life history traits define the timing, intensity, frequency, and duration of key demographic processes along the life cycle of any organism^7,8^. In life-history PCAs, life history traits are reduced to a series of dominant axes that capture the dominant combinations of life history traits and their trade-offs, thus offering a quantitative perspective on the emerging life history strategies (e.g.,^72^). In plants and animals, PCAs of life history traits have highlighted the existence of two dominant axes reflecting a trade-off between development and survival (the fast-slow continuum^73,74^, and the different reproduction strategies, from semelparous to extremely iteroparous (the reproductive strategies continuum^74^.

Here, we derived the following six key life history traits from our examined MPMs: (1) individual development (*γ*); (2) mean life expectancy (*η*_*e*_), (3) distribution of mortality risk along the life cycle (*P*); (4) probability of achieving reproduction before dying (*p*_*α*_); (5) mean age at first reproduction (*L*_*a*_); and (6) reproductive window (*L*). The combination of these six life history traits in populations of animals or plants adequately defines their life history strategies via multivariate analyses such as principal component analyses (PCA^12,14,73^). Once the life history traits were derived, we ensured a robust implementation of the PCA by removing outlier populations based on the Mahalanobis distance using the *malahanobis_distance* function from the *rstatix* R package^75^. Briefly, this approach calculates the distance of each species population from the centre of the multivariate distribution and removes those populations whose distance is greater than expected by chance, assuming a 99% C.I (Table S2).

We performed a PCA using the *prcomp* function from the *base* R package. We retained the first two principal components (PC) because they together captured a similar amount of variation to that achieved by recent studies used to define the fast-slow and reproductive strategy continua^14,74^. Together, these *PC*1_*LH*_ (life history PC1) and *PC*2_*LH*_ explained 66.4% of the variation in life history traits (*PC*1_*LH*_: 38.0%; *PC*2_*LH*_: 28.4%; Figure S1). To quantify the position of each population in the two-dimensional space defined by *PC*1_*LH*_ and *PC*2_*LH*_, we obtained the PCA scores with the function *scores* of the *Vegan* R package v2.5.7^76^.

### Environmental data, environmental stochasticity, and the environmental PCA

To test *H*_*3*_, if higher stochasticity in environmental variables pushes less buffered responses, we linked demographic responses to environmental stochasticity. We started by quantifying the environmental stochasticity experienced by our 134 examined populations, and extracted environmental information from CHELSAcruts^60^. To do so, we downloaded 1 km^2^ gridded monthly values of maximum and minimum temperature and total monthly precipitation that corresponded to latitude and longitude, as well as the years of each study selected from COMPADRE and COMADRE. The CHELSAcruts database includes climatic information from 1901 to 2016, thus including the temporal extent of all populations used in this study (earliest start in 1938 and latest end in 2011; See supplementary material).

Because we are interested in demographic responses to environmental stochasticity rather than mean temperature or precipitation, we decomposed each climatic variable *Y* into three groups of variables that represent trend (*T*), seasonality (*S*), and stochasticity (*R*). We derived these variables using a seasonal decomposition through an additive moving averages model toward the *decompose* function in the *base* R package^77^. Briefly, this function computes the trend component of the variable of interest (*e*.*g*., monthly temperature) using an additive moving average approach. This trend is then subtracted from the time series. The resulting time series represents the periodic fluctuation - the seasonality component (*S*). Finally, trend and season are discounted from the raw time series and the residuals represent the stochasticity of the environmental component - *R* ^77^. For each component of time series *Y* (*i*.*e*., *T*_*Y*_, *S*_*Y*_, *R*_*Y*_), this approach allow us to derivate the mean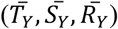, relative amplitude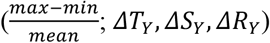, and coefficient of variation (relative stochasticity hereafter,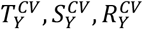). However, we only retained those variables strictly related to change in mean-variance environment and with direct biological or climate meaning. Thus, only the mean trend of the environmental conditions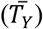, amplitude of trend component (*ΔT*_*R*_) and the relative stochasticity of the random component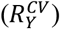. While the later component represents the monthly environmental uncertainty, the other environmental variables represent how environment has changed during the study time compared to the mean conditions.

Similar to the life history PCA described above, we also applied a PCA to environmental data (Environmental PCA, *PC*_*Env*_ hereafter) to retain those variables that best describe variation in environmental conditions across our 134 populations. Before performing the PCA, we assessed the collinearity of the environmental variables by reducing the highly correlated (|ρ| > 0.8 as the threshold; Figure S2). As such, only the following six variables were further examined through our environmental PCA: (1) Mean value of the maximum temperature trend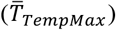, (2) relative stochasticity of maximum temperature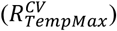, (3) relative amplitude of the trend of the maximum temperature (*ΔT*_*TempMax*_), (4) relative amplitude of the trend of the minimum temperature (*ΔT*_*TempMin*_), (5) mean value of the precipitation trend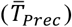, and (6) relative amplitude of the trend of the maximum temperature (*ΔT*_*prec*_). The first two PCs emerging from this next step explained 50.9% of the variation in environmental stochasticity separating those populations from lower to higher stochastic environments (Figure S2). Principal component axis 1 (*PC*1_*Env*_) explains 30.6% of the variation, and describes relative monthly stochasticity in temperature, 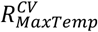, with a gradient from the tropical region to higher latitudes (Figure S3). In contrast, *PC*2_*Env*_, which explains 20.3% of the environmental variation, describes differences in the amplitude of trend in precipitation, *ΔT*_*precipitation*_. *PC*2_*Env*_ is higher in more arid places such as deserts and the Mediterranean region. Therefore, we refer to these environmental axes as temperature stochasticity (*PC*1_*Env*_) and seasonal aridity (*PC*2_*Env*_).

### Statistical analysis and the phylogenetic component

Hypotheses *H*_*1b*_, *H*_*2*_ and *H*_*3*_ were assessed using a single mathematical model (Equation 1) where competing explanatory variables representing life history (*PC*_*LH*_) and environmental stochasticity (*PC*_*Env*_) were used to explain sum of stochastic elasticities with respect to the variance, 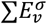.

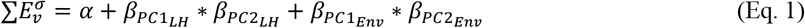

Equation 1 assumes that the capacity of a population to buffer against to environmental stochasticity, 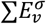, can be predicted by the position of the populations within the two main axes of life histories (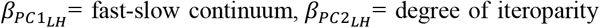; Fig. 1A) and the environmental conditions that the populations faced during the study time (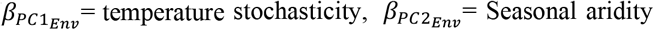; Fig. 1B), including an intercept. The life history axes *PC*1_*LH*_ and *PC*2_*LH*_, along with the environmental axes *PC*1_*Env*_ and *PC*2_*Env*_, were *z*-transformed (*μ* = 0, and sd=1). This normalisation allows for the comparison among *β* estimates across all variables regarding their relative contribution to determine population’s position along the demographic buffering continuum. Here, we further distinguish between *βλ* and *β* to refer to the slope estimates in Eq. 1 with and without phylogenetic corrections, respectively.

We used a Bayesian Phylogenetic Generalised linear model to estimate the most suitable contribution of each of our variables to predict the degree of demographic buffering, 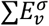. The Bayesian Phylogenetic Generalised linear model was performed via the *MCMCglmm* R package^78^, according to de Villemereuil and Nakagawa^79^. The GLMMs were settled with a flat prior distribution *N*(0,0.2) with 50,000 iterations. Finally, 95% of the posterior density distributions for each variable were compared and accepted as significant when they did not overlap 0 (fake p-value in the MCMCglmm output,^78^). Finally, we estimated the phylogenetic signal using the Pagel’s *λ* (not to be confused with [stochastic] population growth rate *λ* [*λ*_*s*_]), according de Villemereuil and Nakagawa^79^. Briefly, Pagel’s *λ* ranges between 0 and 1, where values close to 0 represent complete randomness in the examined variable and values close to 1 suggest a strong influence of evolutionary history determining the state of the variable^80,81^.

## Supporting information

SOM

## Acknowledgements

This study was primarily financed by CAPES and NERC. GSS was supported by CAPES and CNPq (301343/2023-3). MK was supported by the European Commission through the Marie Skłodowska-Curie fellowship (MSCA MaxPersist #101032484) hosted by RS-G. RS-G was supported by a NERC IRF (NE/M018458/1) and a NERC Pushing the Frontiers (NE/X013766/1).

