## Supplementary material for "Population responses to environmental stochasticity are primarily driven by survival-reproduction trade-offs and mediated by aridity": SOM

This section provides the supplementary information for the manuscript "*Population responses to environmental stochasticity are primarily driven by survival-reproduction trade-offs and mediated by aridity*". Results for life history and environmental Principal Component Analyses are provided in the figures S1 and S2. Populations and their position along the gradient of environmental stochasticity is presented in the figure S3. The proportional contribution of variation in each vital rate to the overall effect of vital rates variation on the stochastic population growth rate $\lambda_s$ is presented in the table S1. Finally, the metadata of the studied populations are presented in the table

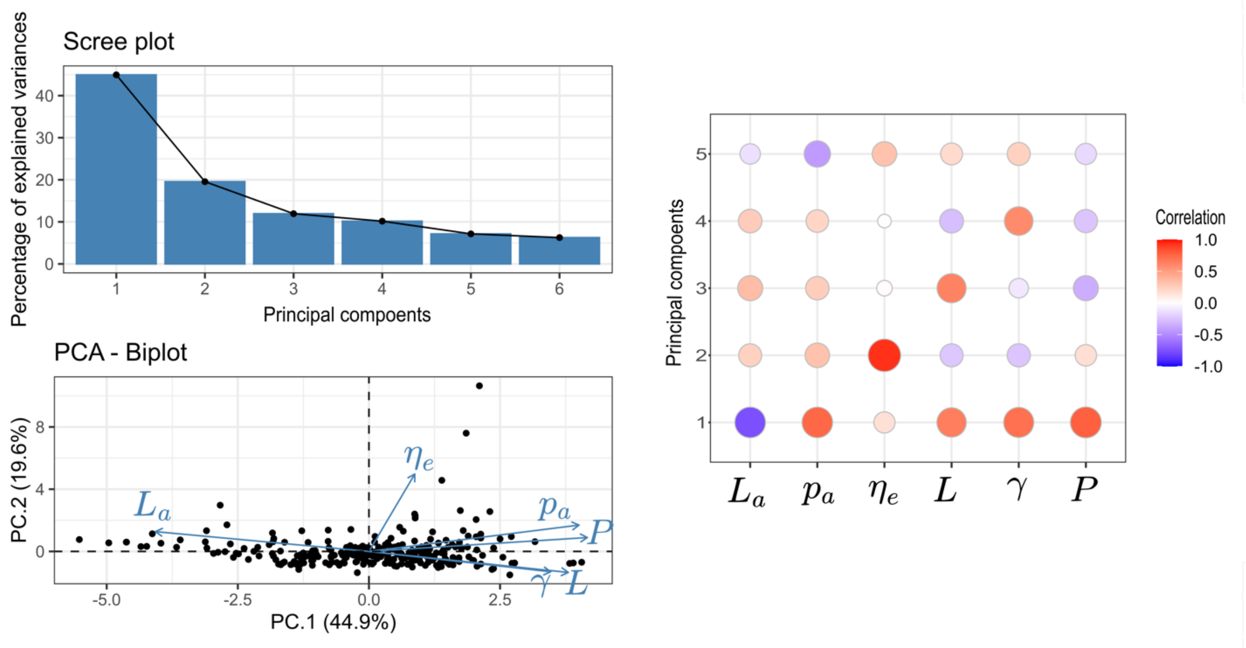


Figure S1. Results from the the life history principal component analysis (PCA) are depicted, showing that the first two principal components are better represented by the stochasticity in temperature and precipitation, respectively. The panel figure shows the scree plot, biplot and correlation plot related to the principal components and the environmental variables. Top right, the scree plot supports that the first two principal components (PCs) capture most of the variance (44.9% and 19.3%, respectively), suggesting that they represent the most important patterns in the pace-of-life. Bottom left, the biplot depicts how the 134 studies populations (13 animals and 121 plants) are distributed along to the environmental gradients captured by the PCs. The arrows represent the life history traits, their lengths indicate their relative contribution to each PC. Notations represent six key life history traits: individual development ($\gamma$); mean life expectancy ($\eta_e$), distribution of mortality risk along the life cycle ($P$); probability of achieving reproduction before dying ($p_a$); mean age at first reproduction ($L_a$); and reproductive window ($L$). PC1 is stronger associated with all aspects of the life history traits, representing how individuals allocate their energy between survival and reproduce – the fast-slow continuum. On the other hand, PC2 is strongly related life expectancy where longer life-spans are related to continuous investment in reproduction (iteroparity) rather than a single reproductive event (semelparity).

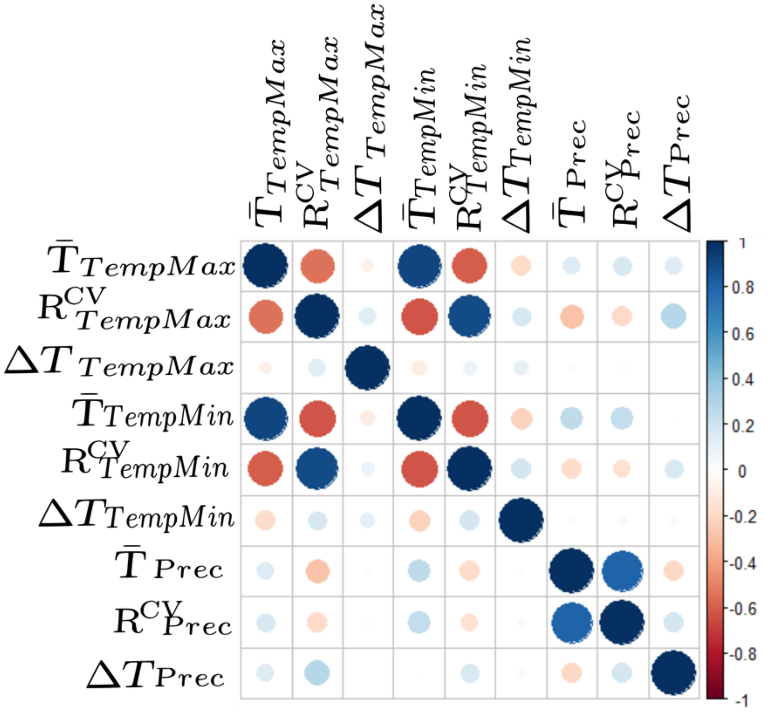


Figure S2. Correlogram showing pairwise correlation used to identify multicollinearity in environmental variables. Notations represent: mean trend ($\bar{\boldsymbol{T}}$), amplitude of the trend ($\boldsymbol{\Delta T}$) and proportional environmental stochasticity ($\boldsymbol{R}^{CV}$). Colours and size represent the person's correlation (ρ). When variables are highly correlated with others (|ρ| > 0.8), only one pair was used in the environmental PCA (see figure S3).

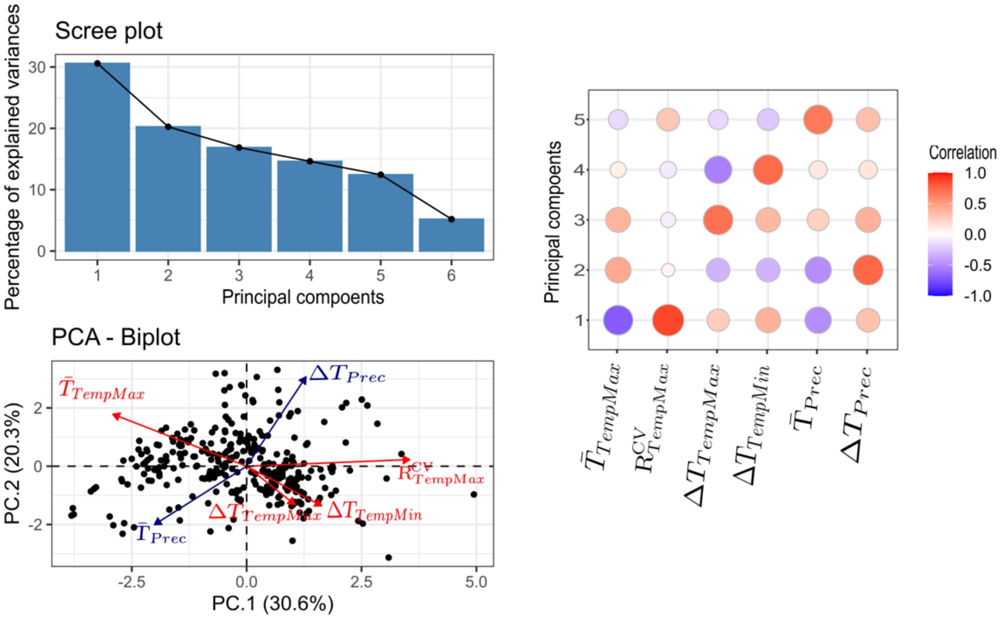


Figure S3. Results from the Environmental Principal Component Analysis (PCA) are depicted, showing that the first two principal components are better represented by the stochasticity in temperature and precipitation, respectively. The panel figure shows the scree plot, biplot and correlation plot related to the principal components and the environmental variables. Top right, the scree plot supports that the first two principal components (PCs) capture most of the variance (30.6% and 20.3%, respectively), suggesting that they represent the most important patterns in the environmental data, but there is still a large amount of variation to be explored by further research. Bottom left, the biplot depicts how the 134 studies populations (13 animals and 121 plants) are distributed along to the environmental gradients captured by the PCs. The arrows represent the environmental variables, their lengths indicate their relative contribution to each PC and their colours are set to blue for variables related to precipitation and red for variables related to temperature. Notations represent: Mean value of the maximum temperature trend ($\boldsymbol{\overline{T}_{TempMax}}$), relative stochasticity of maximum temperature ($\boldsymbol{R^{CV}_{TempMax}}$), relative amplitude of the trend of the maximum temperature ($\boldsymbol{\Delta T_{TempMax}}$), relative amplitude of the trend of the minimum temperature ($\boldsymbol{\Delta T_{TempMin}}$), mean value of the precipitation trend ($\boldsymbol{\overline{T}_{Prec}}$), and relative amplitude of the trend of the maximum temperature ($\boldsymbol{\Delta T_{Prec}}$). PC1 is stronger associated with stochasticity in maximum temperature, while PC2 is mainly determined by the variation in precipitation, as can be better observed in the correlation plot on the right side of the panel.

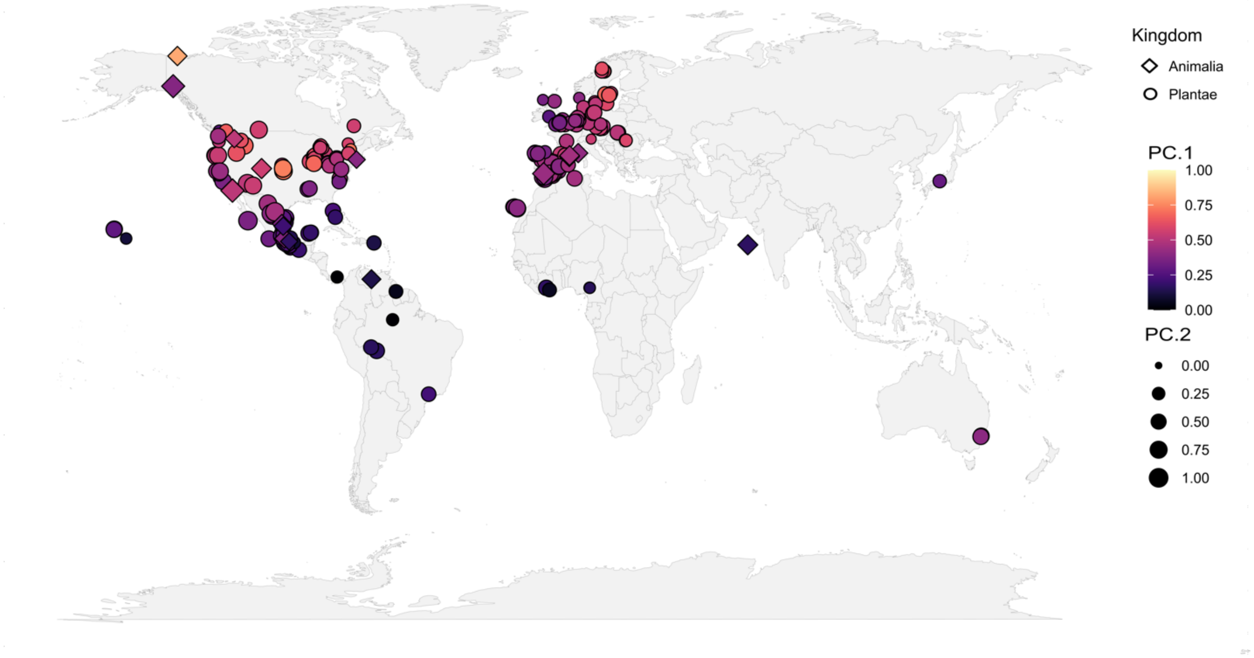

Figure S4. The 134 studied populations (13 animals and 121 plants) are mapped as long their distribution along the environmental gradients captured by the environmental principal components analysis. PC1, representing stochasticity in temperature fluctuations, shows a visual trend to increase with latitude. Despite a more elusive trend, it is possible to identify that PC2 increases from continental to insular areas, however, this pattern might be masked by topographic features (e.g. altitude) which are not shown. Animals and plants are depicted by different shapes – circle for plants and diamonds for animals, and PC's score were proportionally transformed to range from 0 to 1 improving comparison between size and colours.

Table S1. Percentage of contribution of variation in each vital rate $v$, $E_v^{\sigma}$, to the overall effect of temporal variation in on the stochastic population growth rate $\lambda_s$, which is represented by $\sum E_v^{\sigma}$. Animals are grouped by order and plants are grouped by life-forms. Life forms follows COMPADRE v6.23.5.0. The number of populations analysed is denoted by N.

| KINGDOM | TAXA | N | $E_{Survival}^{\sigma}$ | $E_{Growth}^{\sigma}$ | $E_{Reproduction}^{\sigma}$ | $E_{Shrinkage}^{\sigma}$ | $E_{Clonality}^{\sigma}$ |
|---|---|---|---|---|---|---|---|
| **Animalia** | Actinopterygii | 2 | 67.3 | 0.88 | 31.8 | 0 | 0 |
| **Animalia** | Anthozoa | 1 | 54.9 | 1.44 | 1.99 | 41.7 | 0 |
| **Animalia** | Aves | 3 | 82.7 | 0 | 17.3 | 0 | 0 |
| **Animalia** | Insecta | 2 | 39.9 | 0 | 60.1 | 0 | 0 |
| **Animalia** | Mammalia | 1 | 92.5 | 4.11 | 0.74 | 2.69 | 0 |
| **Animalia** | Reptilia | 4 | 69.9 | 22.3 | 7.88 | 0 | 0 |
| **Plantae** | Annual | 4 | 9.32 | 0.04 | 84.5 | 6.14 | 0 |
| **Plantae** | Epiphyte | 1 | 56.6 | 1.77 | 39.7 | 1.94 | 0 |
| **Plantae** | Herbaceous perennial | 92 | 48.9 | 20.7 | 18.3 | 5.73 | 6.30 |
| **Plantae** | Palm | 3 | 41.8 | 24.4 | 21.8 | 10.9 | 1.05 |
| **Plantae** | Shrub | 3 | 71.5 | 27.7 | 0.37 | 0.48 | 0 |
| **Plantae** | Succulent | 10 | 60.9 | 26.7 | 4.95 | 7.50 | 0 |
| **Plantae** | Tree | 8 | 67.2 | 28.0 | 1.84 | 2.91 | 0 |

stochastic elasticities within respect to the variance $\sum E_v^{\sigma}$ is presented for each population.

| | | | | | | | | |
|---|---|---|---|---|---|---|---|---|
| Animalia | *Ursus maritimus* | Mammalia | 183 | 5 | 2001 | 2006 | 69.5000 | -137.0000 |
| Animalia | *Falco naumanni* | Aves | 170 | 5 | 1988 | 1993 | 37.3833 | -5.9667 |
| Animalia | *Lagopus leucura* | Aves | 394 | 5 | 2004 | 2006 | 61.2667 | -138.5167 |
| Animalia | *Lagopus muta* | Aves | 394 | 6 | 2004 | 2006 | 61.2667 | -138.5167 |
| Animalia | *Sceloporus grammicus* | Reptilia | 252 | 5 | 1991 | 1996 | 19.6833 | -98.7000 |

| | | | | | | | | |
|---|---|---|---|---|---|---|---|---|
| Animalia | *Xenosaurus platyceps* | Reptilia | 320 | 3 | 2000 | 2003 | 23.6667 | -99.6333 |
| Animalia | *Xenosaurus platyceps* | Reptilia | 320 | 3 | 2000 | 2003 | 23.0333 | -99.1500 |
| Animalia | *Xenosaurus grandis* | Reptilia | 414 | 4 | 2000 | 2004 | 18.8667 | -97.0167 |
| Animalia | *Clinocottus analis* | Actinopterygii | 87 | 9 | 1996 | 2000 | 32.8167 | -117.3000 |

| | | | | | | | | |
|---|---|---|---|---|---|---|---|---|
| Animalia | *Clinocottus analis* | Actinopterygii | 87 | 11 | 1996 | 2000 | 32.7500 | -117.3000 |
| Animalia | *Colias alexandra* | Insecta | 157 | 3 | 1975 | 1979 | 38.7806 | -106.8900 |
| Animalia | *Scolytus ventralis* | Insecta | 22 | 3 | 1966 | 1968 | 46.9400 | -116.6453 |
| Animalia | *Paramuricea clavata* | Anthozoa | 231 | 4 | 1999 | 2004 | 43.0042 | 6.3961 |
| Plantae | *Euphorbia fontqueriana* | Succulent | 267 | 4 | 2001 | 2004 | 39.8000 | 2.8000 |
| Plantae | *Hypericum cumulicola* | Herbaceous perennial | 463 | 53 | 1994 | 1999 | 27.1667 | -81.3500 |

| | | | | | | | | |
|---|---|---|---|---|---|---|---|---|
| | | perennial | | | | | | |
| Plantae | *Purshia subintegra* | Shrub | 362 | 7 | 1996 | 2003 | 34.7333 | -111.9833 |
| Plantae | *Cucurbita pepo* | Annual | 457 | 3 | 2004 | 2007 | 33.1333 | -90.3500 |
| Plantae | *Castanea dentata* | Tree | 134 | 4 | 1996 | 2000 | 44.4328 | -85.1564 |
| Plantae | *Castanea dentata* | Tree | 134 | 4 | 1996 | 2000 | 45.0525 | -85.7594 |

| | | | | | | | | |
|---|---|---|---|---|---|---|---|---|
| Plantae | *Castanea dentata* | Tree | 134 | 4 | 1996 | 2000 | 44.6436 | -86.2300 |
| Plantae | *Castanea dentata* | Tree | 134 | 4 | 1996 | 2000 | 44.5167 | -86.1069 |
| Plantae | *Castanea dentata* | Tree | 134 | 4 | 1996 | 2000 | 44.4811 | -85.8633 |
| Plantae | *Astragalus scaphoides* | Herbaceous perennial | 326 | 5 | 1986 | 1993 | 45.0667 | -113.0167 |
| Plantae | *Astragalus alopecurus* | Herbaceous perennial | 409 | 6 | 1996 | 2004 | 46.2275 | 2.2136 |
| Plantae | *Astragalus tremolsianus* | Herbaceous perennial | 267 | 3 | 2001 | 2004 | 36.8667 | -2.7667 |
| Plantae | *Dorycnium spectabile* | Herbaceous perennial | 267 | 4 | 2001 | 2004 | 38.1167 | -1.9000 |

| | | | | | | | | |
|---|---|---|---|---|---|---|---|---|
| Plantae | *Anthyllis vulneraria* | perennial | 355 | 4 | 2004 | 2006 | 46.8167 | 11.0333 |
| Plantae | *Lupinus tidestromii* | Herbaceous perennial | 132 | 5 | 2005 | 2007 | 38.1086 | -122.9567 |
| Plantae | *Lupinus tidestromii* | Herbaceous perennial | 132 | 6 | 2005 | 2007 | 38.0853 | -122.9683 |
| Plantae | *Lupinus lepidus* | Herbaceous perennial | 55 | 17 | 1991 | 1995 | 46.2456 | -122.1844 |
| Plantae | *Carapa guianensis* | Tree | 123 | 5 | 2005 | 2009 | -10.0244 | -67.7053 |
| Plantae | *Acer saccharum* | Tree | 330 | 4 | 1951 | 2001 | 40.1500 | -88.1667 |

| | | | | | | | | |
|---|---|---|---|---|---|---|---|---|
| Plantae | *Alliaria petiolata* | perennial | 185 | 3 | 2005 | 2008 | 41.9667 | -85.9833 |
| Plantae | *Alliaria petiolata* | Herbaceous perennial | 185 | 3 | 2005 | 2008 | 40.6833 | -89.4833 |
| Plantae | *Alliaria petiolata* | Herbaceous perennial | 185 | 3 | 2005 | 2008 | 42.7667 | -86.2000 |
| Plantae | *Alliaria petiolata* | Herbaceous perennial | 185 | 3 | 2005 | 2008 | 40.0667 | -88.2000 |
| Plantae | *Alliaria petiolata* | Herbaceous perennial | 185 | 3 | 2005 | 2008 | 41.9667 | -83.9167 |
| Plantae | *Alliaria petiolata* | Herbaceous perennial | 185 | 3 | 2005 | 2008 | 42.9167 | -85.7667 |

| | | | | | | | | |
|---|---|---|---|---|---|---|---|---|
| Plantae | *Sphaeralcea coccinea* | perennial | 129 | 26 | 1938 | 1972 | 38.8000 | -99.2000 |
| Plantae | *Helianthemum polygonoides* | Herbaceous perennial | 267 | 5 | 2001 | 2004 | 40.9500 | -5.4000 |
| Plantae | *Helianthemum teneriffae* | Herbaceous perennial | 267 | 3 | 2001 | 2004 | 28.2833 | -16.4333 |
| Plantae | *Erodium paularense* | Herbaceous perennial | 267 | 5 | 2001 | 2004 | 40.9167 | -3.8167 |
| Plantae | *Erodium paularense* | Herbaceous perennial | 267 | 5 | 2001 | 2004 | 40.4500 | -2.2333 |
| Plantae | *Oenothera deltoides* | Herbaceous perennial | 575 | 12 | 1997 | 2000 | 38.0131 | -121.7917 |

| | | | | | | | | |
|---|---|---|---|---|---|---|---|---|
| Plantae | *Plantago coronopus* | Herbaceous perennial | 608 | 3 | 2007 | 2010 | 36.0142 | -5.6044 |
| Plantae | *Plantago coronopus* | Herbaceous perennial | 608 | 3 | 2007 | 2010 | 42.5733 | -9.0725 |
| Plantae | *Plantago coronopus* | Herbaceous perennial | 608 | 3 | 2007 | 2010 | 56.4236 | 12.6272 |
| Plantae | *Dracocephalum austriacum* | Herbaceous perennial | 151 | 3 | 2003 | 2006 | 49.9333 | 14.1833 |
| Plantae | *Dracocephalum austriacum* | Herbaceous perennial | 151 | 3 | 2003 | 2006 | 49.9333 | 14.1333 |

| | | | | | | | | |
|---|---|---|---|---|---|---|---|---|
| | | perennial | | | | | | |
| Plantae | *Ramonda myconi* | Herbaceous perennial | 445 | 6 | 1992 | 1997 | 42.3167 | 1.7833 |
| Plantae | *Myosotis ramosissima* | Annual | 152 | 3 | 2000 | 2002 | 50.5167 | 13.9833 |
| Plantae | *Gentiana pneumonanthe* | Herbaceous perennial | 423 | 3 | 1987 | 1991 | 52.8106 | 6.3925 |
| Plantae | *Hypochaeris radicata* | Herbaceous perennial | 282 | 3 | 1999 | 2003 | 52.0167 | 6.5500 |
| Plantae | *Echinacea angustifolia* | Herbaceous perennial | 129 | 30 | 1938 | 1972 | 38.1333 | -99.0500 |
| [illegible] | [illegible] | Herbaceous | [illegible] | [illegible] | [illegible] | [illegible] | [illegible] | [illegible] |

| | | | | | | | | |
|---|---|---|---|---|---|---|---|---|
| Plantae | *Ratibida columnifera* | Herbaceous perennial | 129 | 14 | 1938 | 1972 | 38.8000 | -99.2000 |
| Plantae | *Liatris scariosa* | Herbaceous perennial | 166 | 5 | 1998 | 2003 | 41.5167 | -87.9000 |
| Plantae | *Artemisia genipi* | Herbaceous perennial | 355 | 3 | 2004 | 2006 | 46.8167 | 11.0333 |
| Plantae | *Solidago mollis* | Herbaceous perennial | 129 | 19 | 1938 | 1972 | 38.8000 | -99.2000 |
| Plantae | *Cirsium palustre* | Herbaceous perennial | 474 | 3 | 2002 | 2005 | 58.9500 | 17.6000 |
| Plantae | *Cirsium dissectum* | Herbaceous perennial | 283 | 3 | 1999 | 2003 | 52.0000 | 5.6000 |

| | | | | | | | | |
|---|---|---|---|---|---|---|---|---|
| Plantae | *Cirsium dissectum* | perennial | 283 | 3 | 1999 | 2003 | 52.0333 | 6.4333 |
| Plantae | *Cirsium dissectum* | Herbaceous perennial | 283 | 3 | 1999 | 2003 | 52.6000 | 6.1333 |
| Plantae | *Cirsium pitcheri* | Herbaceous perennial | 41 | 4 | 1995 | 2000 | 41.6167 | -87.2000 |
| Plantae | *Cirsium pitcheri* | Herbaceous perennial | 166 | 4 | 1998 | 2003 | 42.4167 | -87.8000 |
| Plantae | *Cirsium undulatum* | Herbaceous perennial | 129 | 25 | 1938 | 1972 | 38.1333 | -99.0500 |

| | | | | | | | | |
|---|---|---|---|---|---|---|---|---|
| Plantae | *Jurinea fontqueri* | perennial | 267 | 4 | 2001 | 2004 | 37.7500 | -3.5500 |
| Plantae | *Centaurea jacea* | Herbaceous perennial | 282 | 4 | 1999 | 2003 | 52.0000 | 5.6000 |
| Plantae | *Centaurea jacea* | Herbaceous perennial | 282 | 4 | 1999 | 2003 | 52.0333 | 6.4333 |
| Plantae | *Cheirolophus metlesicsii* | Herbaceous perennial | 267 | 3 | 2001 | 2004 | 28.3833 | -16.4167 |
| Plantae | *Carum carvi* | Herbaceous perennial | 301 | 3 | 2003 | 2006 | 65.3592 | 15.7358 |
| Plantae | *Carum carvi* | Herbaceous perennial | 301 | 3 | 2003 | 2006 | 65.3389 | 15.7142 |

| | | | | | | | | |
|---|---|---|---|---|---|---|---|---|
| Plantae | *Succisa pratensis* | perennial | 282 | 4 | 1999 | 2003 | 52.2833 | 5.7333 |
| Plantae | *Succisa pratensis* | Herbaceous perennial | 282 | 3 | 1999 | 2003 | 52.0333 | 6.4333 |
| Plantae | *Succisa pratensis* | Herbaceous perennial | 282 | 4 | 1999 | 2003 | 52.0000 | 5.6000 |
| Plantae | *Succisa pratensis* | Herbaceous perennial | 381 | 15 | 2000 | 2003 | 58.8333 | 17.4000 |
| Plantae | *Ardisia escallonioides* | Shrub | 431 | 3 | 1992 | 1994 | 25.5000 | -80.5000 |
| Plantae | *Androsace elongata* | Annual | 152 | 5 | 2000 | 2002 | 50.5167 | 13.9833 |
| Plantae | *Primula farinosa* | Herbaceous perennial | 582 | 3 | 2000 | 2006 | 56.5850 | 16.5217 |

| | | | | | | | | |
|---|---|---|---|---|---|---|---|---|
| Plantae | *Primula veris* | Herbaceous perennial | 73 | 12 | 1999 | 2003 | 50.7500 | 5.7500 |
| Plantae | *Primula elatior* | Herbaceous perennial | 272 | 5 | 2004 | 2007 | 50.8500 | 4.8833 |
| Plantae | *Primula vulgaris* | Herbaceous perennial | 596 | 10 | 1992 | 1994 | 51.7778 | 0.6939 |
| Plantae | *Primula vulgaris* | Herbaceous perennial | 170 | 14 | 1999 | 2002 | 51.1833 | 3.3500 |
| Plantae | *Primula vulgaris* | Herbaceous perennial | 591 | 24 | 2008 | 2010 | 43.2833 | -5.5000 |

| | | | | | | | | |
|---|---|---|---|---|---|---|---|---|
| Plantae | *Paronychia jamesii* | Herbaceous perennial | 129 | 25 | 1938 | 1972 | 38.1333 | -99.0500 |
| Plantae | *Atriplex canescens* | Shrub | 603 | 3 | 1995 | 1998 | 26.5083 | -103.5083 |
| Plantae | *Neobuxbaumia mezcalaensis* | Succulent | 182 | 3 | 1999 | 2002 | 18.3333 | -97.4667 |
| Plantae | *Neobuxbaumia mezcalaensis* | Succulent | 183 | 3 | 1999 | 2002 | 18.3333 | -97.4667 |
| Plantae | *Neobuxbaumia macrocephala* | Succulent | 183 | 3 | 1999 | 2002 | 18.3333 | -97.4667 |
| Plantae | *Mammillaria huitzilopochtli* | Succulent | 193 | 5 | 1999 | 2004 | 17.8114 | -96.9639 |

| | | | | | | | | |
|---|---|---|---|---|---|---|---|---|
| Plantae | *Mammillaria solisioides* | Succulent | 490 | 3 | 1999 | 2002 | 18.0831 | -97.9178 |
| Plantae | *Ariocarpus fissuratus* | Succulent | 350 | 4 | 2005 | 2009 | 26.9119 | -102.1225 |
| Plantae | *Ariocarpus fissuratus* | Succulent | 350 | 3 | 2005 | 2009 | 26.9111 | -102.1269 |
| Plantae | *Escobaria robbinsorum* | Succulent | 511 | 5 | 1988 | 1993 | 34.1333 | -109.9333 |
| Plantae | *Rumex rupestris* | Herbaceous perennial | 267 | 4 | 2001 | 2004 | 43.0000 | -9.2500 |
| Plantae | *Armeria caespitosa* | Herbaceous perennial | 208 | 24 | 2005 | 2008 | 40.7667 | -3.9333 |

| | | | | | | | | |
|---|---|---|---|---|---|---|---|---|
| Plantae | *Limonium erectum* | perennial | 267 | 5 | 2001 | 2004 | 40.3833 | -2.9000 |
| Plantae | *Ranunculus peltatus* | Herbaceous perennial | 266 | 3 | 1992 | 1997 | 59.1333 | 16.0000 |
| Plantae | *Actaea spicata* | Herbaceous perennial | 204 | 6 | 1993 | 1998 | 58.9500 | 17.6167 |
| Plantae | *Actaea spicata* | Herbaceous perennial | 204 | 5 | 1993 | 1998 | 58.9833 | 17.5500 |
| Plantae | *Dicentra canadensis* | Herbaceous perennial | 331 | 4 | 2008 | 2011 | 40.3514 | -82.9264 |
| Plantae | *Dicentra canadensis* | Herbaceous perennial | 331 | 6 | 2008 | 2011 | 40.1200 | -82.9669 |

| | | | | | | | | |
|---|---|---|---|---|---|---|---|---|
| Plantae | *Dicentra canadensis* | perennial | 331 | 6 | 2008 | 2011 | 39.5861 | -82.5464 |
| Plantae | *Chamaedorea elegans* | Palm | 594 | 6 | 1996 | 1999 | 17.9500 | -96.5000 |
| Plantae | *Chamaedorea radicalis* | Palm | 172 | 14 | 1999 | 2005 | 23.0833 | -99.1667 |
| Plantae | *Borassus aethiopum* | Palm | 36 | 4 | 1996 | 1998 | 6.2167 | -5.0333 |
| Plantae | *Poa alpina* | Herbaceous perennial | 355 | 5 | 2004 | 2006 | 46.8167 | 11.0333 |

| | | | | | | | | |
|---|---|---|---|---|---|---|---|---|
| Plantae | *Zea diploperennis* | Herbaceous perennial | 500 | 11 | 1988 | 1993 | 19.6000 | -104.2667 |
| Plantae | *Danthonia sericea* | Herbaceous perennial | 384 | 3 | 1983 | 1985 | 36.0000 | -78.9333 |
| Plantae | *Hilaria mutica* | Herbaceous perennial | 601 | 4 | 1996 | 1998 | 26.0000 | -103.0000 |
| Plantae | *Catopsis compacta* | Epiphyte | 143 | 3 | 2005 | 2008 | 17.2653 | -96.5617 |

| | | | | | | | | |
|---|---|---|---|---|---|---|---|---|
| Plantae | *Orchis purpurea* | perennial | 273 | 16 | 2002 | 2008 | 50.7500 | 5.7500 |
| Plantae | *Cypripedium fasciculatum* | Herbaceous perennial | 576 | 7 | 1999 | 2007 | 42.3333 | -123.5167 |
| Plantae | *Cypripedium fasciculatum* | Herbaceous perennial | 576 | 7 | 1999 | 2007 | 42.2500 | -123.1667 |
| Plantae | *Cypripedium fasciculatum* | Herbaceous perennial | 576 | 5 | 1999 | 2007 | 42.3333 | -122.4500 |
| Plantae | *Allium tricoccum* | Herbaceous perennial | 402 | 4 | 1984 | 1988 | 45.5167 | -75.9167 |
| Plantae | *Trillium grandiflorum* | Herbaceous perennial | 304 | 5 | 1999 | 2001 | 41.5167 | -80.4333 |
| Plantae | *Trillium grandiflorum* | Herbaceous perennial | 304 | 5 | 1999 | 2001 | 41.7167 | -80.0333 |
| Plantae | *Trillium grandiflorum* | Herbaceous | 304 | 5 | 1999 | 2001 | 41.8000 | -80.2000 |

| | | | | | | | | |
|---|---|---|---|---|---|---|---|---|
| | | perennial | | | | | | |
| Plantae | *Trillium grandiflorum* | Herbaceous perennial | 304 | 5 | 1999 | 2001 | 41.4833 | -80.4000 |
| Plantae | *Trillium grandiflorum* | Herbaceous perennial | 304 | 3 | 1999 | 2001 | 41.3333 | -79.9667 |
| Plantae | *Trillium grandiflorum* | Herbaceous perennial | 304 | 7 | 1999 | 2001 | 41.6000 | -80.3500 |
| Plantae | *Trillium grandiflorum* | Herbaceous perennial | 304 | 3 | 1999 | 2001 | 41.6833 | -80.0667 |
| Plantae | *Trillium grandiflorum* | Herbaceous perennial | 304 | 4 | 1999 | 2001 | 41.7500 | -79.9500 |
| Plantae | *Calochortus lyallii* | Herbaceous perennial | 382 | 32 | 1996 | 2000 | 49.0333 | -119.6167 |
| Plantae | *Asarum canadense* | Herbaceous perennial | 131 | 14 | 1989 | 1995 | 44.0000 | -75.0000 |

| | | | | | | | | |
|---|---|---|---|---|---|---|---|---|
| Plantae | *Pinus strobus* | Tree | 391 | 9 | 2005 | 2007 | 50.8667 | 14.4167 |
